# A robust and transformation-free joint model with matching and regularization for metagenomic trajectory and disease onset

**DOI:** 10.1101/2022.04.19.488854

**Authors:** Qian Li, Kendra Vehik, Cai Li, Eric Triplett, Luiz Roesch, Yi-Juan Hu, Jeffery Krischer

## Abstract

**Background:** To identify operational taxonomy units (OTUs) signaling disease onset in an observational study, a powerful strategy was selecting participants by matched sets and profiling temporal metagenomes, followed by trajectory analysis. Existing trajectory analyses modeled individual OTU or microbial community without adjusting for the within-community correlation and matched-set-specific latent factors.

**Results:** We proposed a joint model with matching and regularization (JMR) to detect OTU-specific compositional trajectory predictive of host disease status, using nested random effects and covariate taxa pre-selected by Bray-Curtis distance and elastic net regression. The inherent negative correlation in microbiota composition was adjusted by incorporating the top-correlated taxa as covariate. We designed a simulation pipeline to generate true biomarkers for disease onset and the pseudo biomarkers caused by compositionality or latent noises. We demonstrated that JMR effectively controlled the false discovery and pseudo biomarkers in a simulation study that generated temporal high-dimensional metagenomic counts with random intercept or slope. Application of the competing methods in the simulated data and the TEDDY cohort showed that JMR outperformed the other methods and identified important taxa in infants’ fecal samples with dynamics preceding host disease status.

**Conclusion:** Our method JMR is a robust framework that models taxon-specific compositional trajectory and host disease status in the matched participants, improving the power of detecting disease-predictive microbial features in certain scenarios.

## Background

Gut microbiota profiled by 16s rRNA gene sequencing or metagenomic (i.e., wholegenome shotgun) sequencing has been frequently used in observational studies of environmental exposures, immune biomarkers, and disease onset [1, 2, 3, 4, 5]. One of the challenges in analyzing microbiota in an observational study is to incorporate the matching between participants by certain confounding risk factors (e.g. gender, clinical site, etc.) and/or disease status (case-control), such as the DIABIMMUNE and TEDDY cohorts [1, 2, 5]. A matching design effectively eliminates the noise effect of sample collection, storage, shipment, sequencing batch, and environmental exposures confounding with disease outcomes, as well as reduces the costs of sample collection and sequencing. Statistical analyses for microbiota in matched sets included, but are not limited to, conditional logistic regression [1], non-parametric comparison PERMANOVA [6] and LDM [7] with extension to compare cases and controls within a matched set [8], which aimed to model and analyze microbiome data at independent time points.

Longitudinal profiling is a powerful strategy for the microbiome studies that aim to identify differential microbial trajectories between exposure groups or phenotypes [9, 10] or detect the time intervals of differential abundance [11]. However, most of these studies failed to test if the compositional trajectory of an operational taxonomic unit (OTU) signaled host disease status. To detect microbial trajectories predictive of disease outcome in matched sets, an intuitive method is the generalized linear mixed effect model with or without the zero-inflation component [9, 10, 12], in which a taxon’s abundance and/or presence is the outcome variable and the disease status is the covariate of interest. The ZIBR model [9] tests the association between OTU and a covariate factor using a two-part model for the non-zero relative abundance and presence of each OTU, assuming the non-zero relative abundance and presence being independent. A similar framework [10] was proposed to analyze the longitudinal zero-inflated counts per OTU using a Negative Binomial distribution, without converting the raw counts to relative abundance. A semi-parametric approach for longitudinal taxon-specific relative abundance is the linear mixed effect model (LMM) with asin-square-root transformation, which has been implemented in MaAsLin 2 [12].

One concern about using generalized linear mixed model to test the association between 16S rRNA or metagenomic trajectory and disease onset is that the covariates in this model may contribute to disease risk. For example, the HLA haplogenotypes and early use of probiotics may affect infants’ gut microbiota and should be included as covariates. These factors were also found associated with islet autoimmunity among children enrolled in TEDDY [13]. One usually added interaction terms between each covariate and the disease outcome [3, 12] to adjust for the association. However, a linear model with many interaction terms may lead to over-fitting and reduce the detection power [14]. A sensible choice is the joint modeling of longitudinal biomarker and survival outcomes [15, 16], but there are limitations in applying this model to microbiome data in observational studies. First, the cost of metagenomic sequencing and the availability of fecal samples in a multi-center study may restrict the metagenome profiling to a subgroup of participants selected by certain criteria [1, 2, 3], whose survival outcome may deviate from common statistical assumptions. Second, the classic joint modeling approach aims to address repeated measurements of biomarkers in a time-to-event analysis rather than to test if a biomarker’s intercept or slope is predictive of host health condition. Third, in an observational study that selects and matches participants by certain factors, their risk of developing disease is also matched. Thus, a survival submodel may not be capable of characterizing the disease risk between matched participants.

Many of the existing methods for microbiome data are built on the transformed relative abundance, such as centered log-ratio or inter-quartile log-ratio. In our new method, transformation of compositional data is not considered, since transformation strategy may have profound impact on analysis result and interpretation [17].

The compositional change in true biomarkers (e.g., causal OTUs contributing to disease onset) always leads to simultaneous change in some other OTUs’ composition because of sum-to-one constraint. In an observational study with matching design, it is common to collect and profile microbiota at many time points. The sum-to-one constraint and latent noise effect on microbiota may yield pseudo biomarkers with relative abundance associated with host disease status but not contributing to disease development. Hence, a taxon-level model is built for relative abundance trajectory that adjusts for the dynamic interdependence between taxa and reduces pseudo biomarker rate. In addition, we illustrate the performance of our method by a simulation pipeline that mimics the negative correlation in microbial community. The latent technical noise in microbiome was removed by converting raw counts to relative abundance, and Zero-Inflated Beta density [9] was adopted to model an OTU’s non-zero relative abundance and presence, respectively. We employed a subject-level random effect to link the logistic regression model of disease to a two-part longitudinal submodel. The latent effect of exposures related to matched set indicator was modeled by another random effect nested with the subject-level random effect. The OTU-disease association was assessed by jointly testing the scaling parameters for the subject-level random effect in the two-part submodel. We benchmarked the robustness and power of our method by a comprehensive simulation study and an application in the TEDDY cohort. The results illustrated that our method controlled the rates of false discovery and pseudo biomarkers, as well as improved the efficacy of detecting microbial trajectories signaling disease outcome.

## Results

For simplicity, the aim of present research is to incorporate the matching of microbiome samples and adjust for the unknown dependence between taxa in univariate trajectory analysis, rather than model the microbiota compositionality. Here we develop a Joint model with Matching and Regularization (JMR) to detect taxon-specific compositional trajectory associated with disease onset, adjusting for the top-correlated taxa and matched-set-specific latent noises. According to the characteristics of disease risk and infant-age gut microbiota in the TEDDY cohort, we designed a simulation pipeline similar to [8], generated the observed counts of temporal microbiota and compared our method to LMM and ZIBR using the simulated data. We also applied these methods to the shotgun metagenomic sequencing data profiled from the 4-9 months stool samples of infants enrolled in TEDDY cohort.

### Overview of TEDDY microbiome study

TEDDY is an observational prospective study of children at increased genetic risk of type 1 diabetes (T1D) conducted in six clinical centers in the U.S. and Europe (Finland, Germany, and Sweden). A total of 8,676 children were enrolled from birth and followed every 3 months for blood sample collection and islet autoantibody measurement up to 4 years of age, then every 3-6 months based on autoantibody status until the age of 15 years or diabetes onset [18]. A primary disease endpoint in TEDDY is islet autoimmunity (IA), defined as persistent positive for insulin autoantibodies (IAA), glutamic acid decarboxylase autoantibodies (GADA), or insulinoma-associated-2 autoantibodies (IA-2A) at two consecutive visits confirmed by the two TEDDY laboratories [18]. The participants’ monthly stool samples were collected from 3-month age until the onset of IA or censoring with random missing samples [1, 2]. Based on the sample availability and metagenomic sequencing cost, the microbiome study in TEDDY selected all the participants (cases) who developed IA by the design cutoff date May 31, 2012 and the controls at 1:1 case-control ratio matched by clinical center, gender, family history of T1D to profile the temporal gut microbiota, resulting in S = 418 matched sets (or pairs [19]). These matching factors are known risk factors for type 1 diabetes. Some of the matched sets are at higher risk of IA than the others due to higher risk human leukocyte antigen (HLA) genotypes, geography or having family history of T1D. Hence, the matched participants have comparable risk of IA, but heterogeneity still exists between them according to the case-control status by the design freeze date. The observed metagenomic counts table in TEDDY was generated by the standard procedure of DNA extraction, PCR amplification, shotgun metagenomic sequencing, assembly, annotation and quantification, as described in [1]. We visualized the top abundant species in the metagenomes of TEDDY participants who had matched IA endpoint no later than 2 years of age (**Figure 1**).

**Figure 1.**
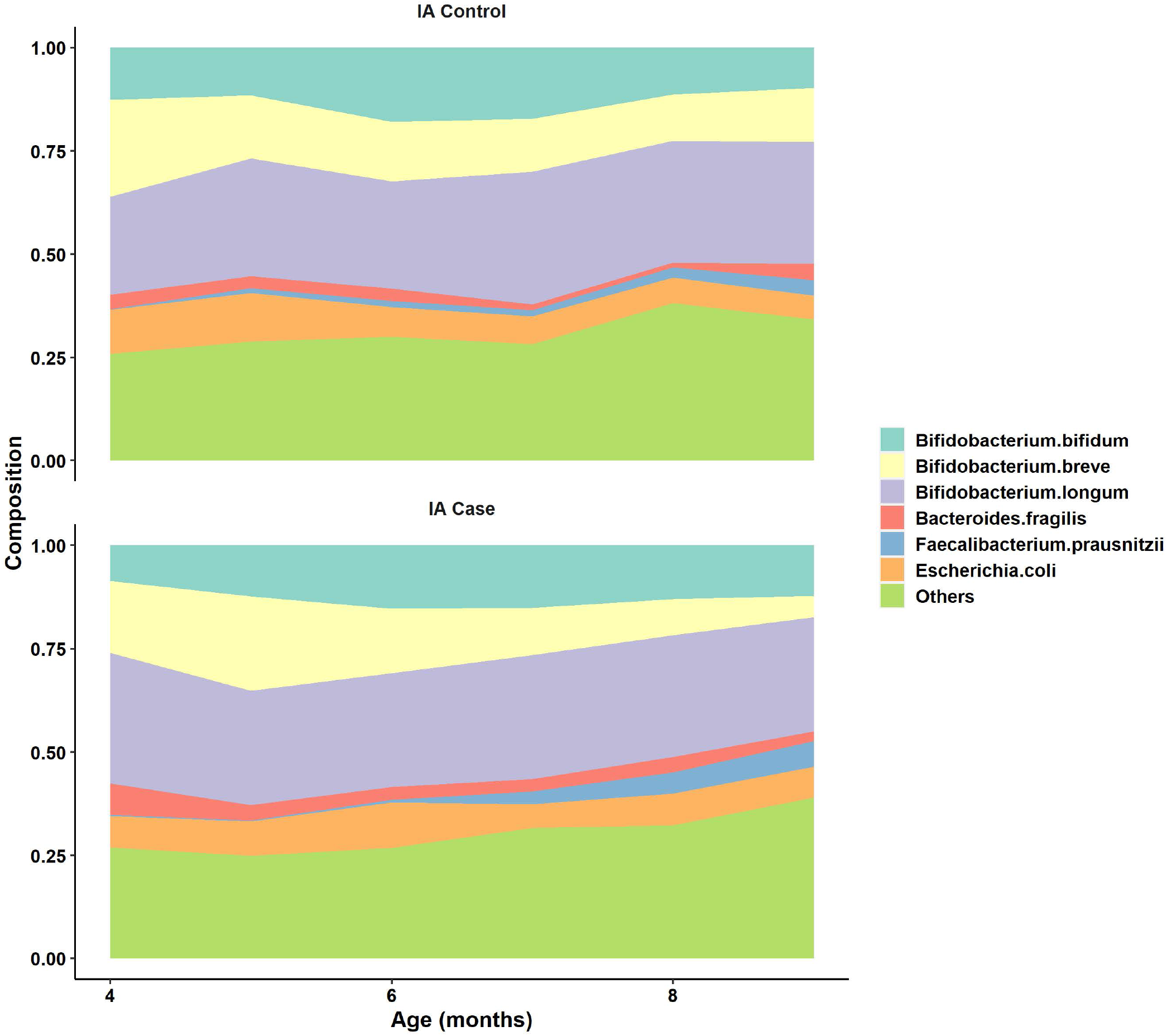
Mean composition trajectory of top abundant species in infant-age metagenomes grouped by host islet autoimmunity status at 2 years of age for a subgroup of TEDDY participants.

### Simulation

Disease outcome for the matched participants are simulated by the procedure below. The observed relative abundance per taxon were simulated by different scenarios. We first generated raw counts for a single OTU by Beta-Binomial distribution to assess the robustness and power of our method JMR without covariate taxa. We also designed a shifting procedure to mimic inherent negative correlation in the true composition of microbiota and generated the temporal high-dimensional raw counts table to evaluate the performance of compared methods.

#### Generate disease outcome in matched sets

We defined matched sets and subjects as ‘high-risk’ and ‘low-risk’ to generate the temporal OTU counts prior to disease onset. Subjects are matched at 1:1 ratio. We first generated subject-level and set-level random effects from a standard Normal distribution 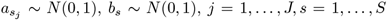. Each random effect was converted to a binary variable by the median value. That is 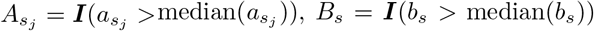, where 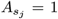 (or *B*_*s*_ = 1) represents a ‘high-risk’ subject (or set). Next, we simulated a genetic risk factor 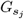 at subject-level with effect size estimated from TEDDY data, and the disease status per subject by a Bernoulli distribution 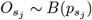, where logit 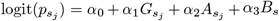. We set (*α*_0_, *α*_1_, *α*_3_) = (0.5, −2, 1) and *α*_2_ ∈ {0.5, 0.75, 1, 1.25, 1.5} to generate different datasets, with *α*_1_ estimated from the real data by JMR.

#### Scenario A: single OTU counts

We first simulated the observed counts of a single OTU by Beta-Binomial [14] distribution to compare the univariate trajectory methods without adjusting for covariate taxa. The true relative abundance of an OTU at the earliest time point *t* = 1 was drawn from a Beta distribution *µ*_1_ ∼ *Beta*(*µ*_0_, *ϕ*_0_), where parameters *µ*_0_, *ϕ*_0_ were estimated by applying Beta-Binomial MLE to the metagenomic raw counts of an OTU selected at a given relative abundance level in the TEDDY data. To simplify the age-dependent effect, the relative abundance of this OTU at later time points *t* > 1 was generated by linearly increasing *µ*_1_ to *µ*_*t*_. The baseline relative abundance at time *t* in a matched set s was generated by *µ*_*st*_ ∼ *Beta*(*µ, ϕ*_*t*_), and was increased or decreased by 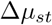 if the set was labeled as ‘high-risk’. The true relative abundance of this OTU for subject *j* in set *s* at time point t was simulated by 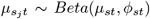, and was increased or decreased by 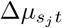 if the subject was ‘high-risk’. The total counts per sample, i.e., library size was drawn from a Poisson distribution 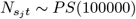, and the counts for this OTU is generated from a Binomial (BN) distribution 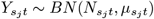.

#### Scenario B1: counts table with random intercept and pseudo biomarkers

We also generated a high-dimensional counts table with *P* = 1030 OTUs to demonstrate the performance of each method, so that the covariate taxa can be used in JMR. First, we simulated the true composition of each microbiome sample by a shifting procedure combined with Dirichlet distribution, generating the negative correlation within microbial community. Next, the sample-wise library size was generated by a Poisson distribution, and the observed raw counts were sampled from a Multinomial distribution, using the aforementioned true composition and sequencing library size. We visualized the simulation procedure of Scenario B1 in **Figure 2** for dimension of *P* = 4.

**Figure 2.**
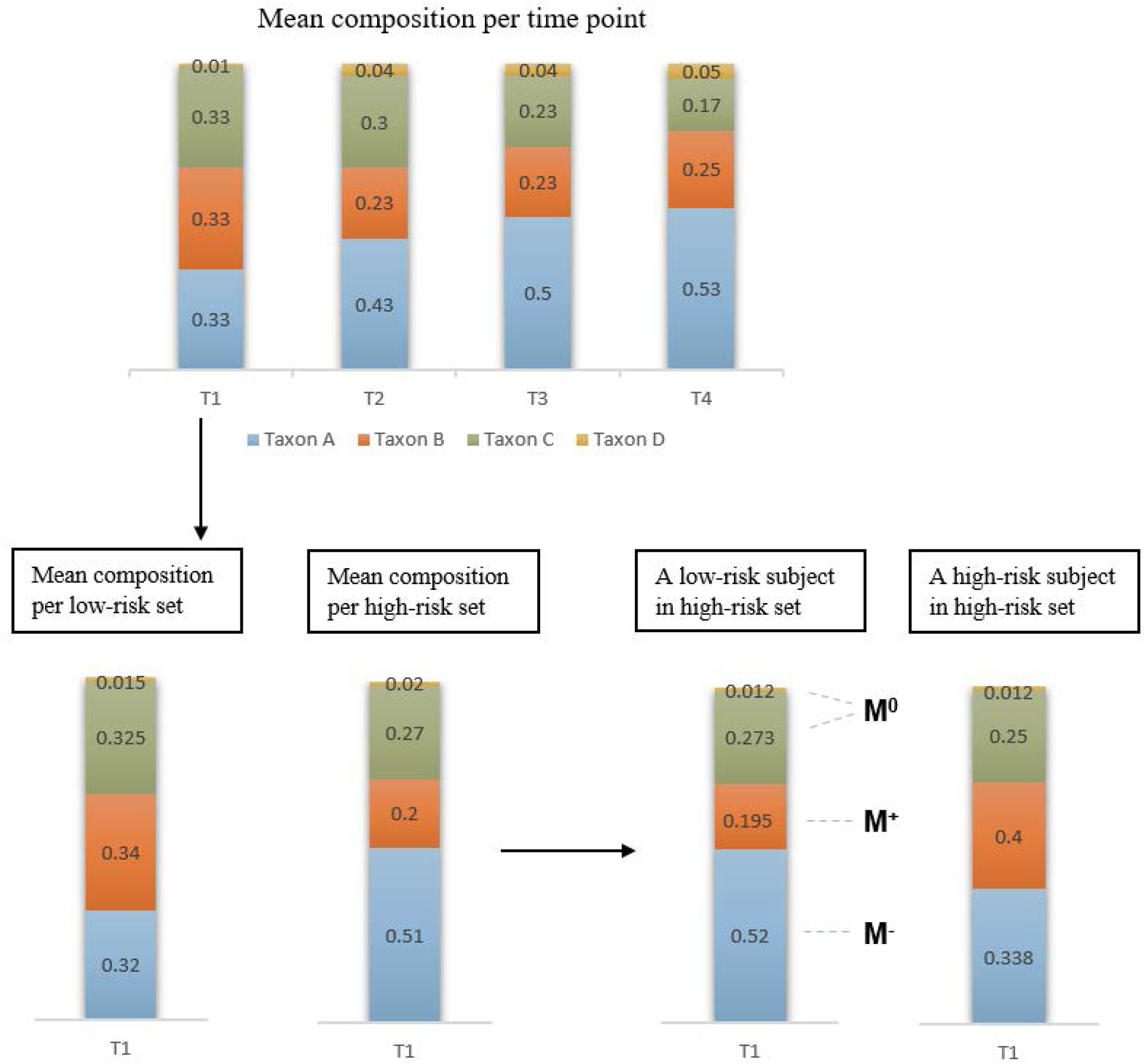
Shifting procedure in simulation scenario B1.

**Step 1**: Estimate the baseline mean composition (or frequency) of microbiota 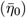 and the overdispersion (*ξ*_0_ = 0.04) at the starting time point *t* = 1 in TEDDY data by Dirichlet-Multinomial (DM) maximum likelihood estimate (MLE) of the observed counts. Generate the mean frequency of microbiota at the first time point by Dirichlet (DL) distribution: 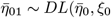.

**Step 2**: The mean frequency 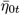 at a later time point *t* > 1 is generated by the following shifting procedure: increase the composition of some OTUs in 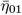 (denoted by 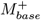) with a sum of Δ_*t*_ and simultaneously reduce the frequency of other OTUs in 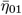 (denoted by 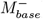) by Δ_*t*_. The absolute shift size Δ_*t*_ represented the age effect on microbiota. This shifting strategy characterized the inherent correlation between 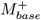 and 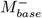 because of the simultaneous compositional change in these OTUs. All the OTUs in 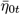 are assigned to either 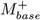 or 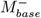 to account for the impact of various latent exposures across time points.

**Step 3**: At each time point, the heterogeneity between matched sets is the overdispersion estimated by DM MLE based on the samples per time point in TEDDY, denoted by *ξ*_*t*_. The overdispersion at the first time point is *ξ*_1_ = 0.05 and linearly decreases over time, which mimics the time-dependent overdispersion observed in the infant-age metagenome in TEDDY. We generated a mean frequency for each matched set *s* at time point *t* by 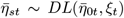. If a set is labeled as ‘high-risk’, we shifted all the OTUs in 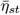 using the procedure in Step 2 with shift size Δ_*st*_, which is a proportion of the maximum shift size, i.e., 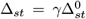. We set *γ* ∈ {0.6, 0.7, 0.8} to evaluate method performance at different set-level random effect size.

**Step 4**: The between-subject heterogeneity within each matched set was the median DM MLE of overdispersion per matched set based on the real data, that is *ξ*^*∗*^ = 0.03. Hence, we generated the true microbiota composition for a sample collected from a ‘low-risk’ subject *j* in set *s* at time *t* by 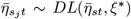. For a subject labeled as ‘high-risk’, we increased 15% OTUs (denoted by *M*^+^) in 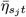 by 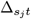, and reduced another 15% OTUs (denoted by *M*^−^) by 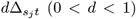. The subsets *M*^+^ and *M*^−^ are the true biomarkers for disease status. We randomly selected a third subset (denoted by *M* ^0^) from the remaining 70% OTUs in 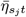 and reduced the composition of *M* ^0^ by a total of 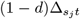. There may exist OTUs never selected in *M*^+^, *M*^−^, or *M* ^0^, which are the ‘null’ OTUs. The OTUs selected in *M* ^0^ are the pseudo biomarkers due to random shift in frequency. Similarly, we set the total shift 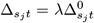 at distinct effect size *λ* ∈ {0.5, 0.6, 0.7, 0.8}, where 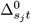 is the maximum shift restricted by sum-to-one.

**Step 5**: The library size for each sample is simulated by a Poisson distribution 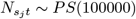, truncated by a minimum of 10000. The raw counts per sample is generated by Multinomial (MN) distribution 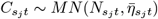.

#### Scenario B2: counts table with random slope and pseudo biomarkers

Data generation process for this scenario is similar to Scenario B1, except that the shift size 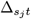 from ‘low-risk’ to ‘high-risk’ subjects varies by time points. It’s worth to note that we cannot distinguish ‘false positive’ from ‘pseudo positive’ in scenarios B1 and B2. Hence, we use the sum of false positive rate and pseudo positive rate, i.e., false or pseudo positive rate (FPPR) as a performance metric for scenarios B1,B2. That is 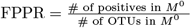.

#### Scenario C: counts table without pseudo biomarkers

In this scenario we considered random intercept signaling the disease onset and fixed half OTUs in 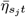 as ‘null ’ in order to evaluate the FPR and FDR of each method, although this scenario is not applicable to real data. Among the other half OTUs, we selected 10% OTUs in 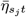 as *M*^+^ and 40% OTUs as *M*^−^ without a subset of pseudo biomarkers (*M* ^0^).

#### Performance of competing methods

In scenario A, we compared JMR not adjusting for correlated taxa (JMR-NC) with the following methods : a) a joint model with regularization but without matching indicator and correlated taxa (JR-NC); b) the ZIBR model with a Wald statistic jointly testing OTU-specific abundance or presence using either a single random effect (ZIBR-S) or nested random effects (ZIBR-N); c) LMM with arcsin-square-root transformation using either a single random effect (LMM-S) or nested random effects (LMM-N). For ZIBR and LMM methods, we used R package gamlss and set the sample age, genotype, disease status, and genotype-disease interaction term as the fixed effect covariates.

We randomly selected 6 OTUs with different relative abundance from TEDDY data and estimated the baseline parameters for each. Then we generated *n* = 10000 replicates for each OTU with *S* ∈ {50, 100}. The type I error rate and power of each method was calculated at statistical significance level *p* < 0.05, shown in Table 1. The results showed that JMR-NC persistently controlled the type I error and provided higher detection power at distinct abundance levels except for the OUTs with relative abundance between (10^−2^, 10^−3^) and (10^−6^, 10^−5^). Type I error of the reduced model JR-NC was severely inflated in some datasets and its power was lower than JMR. LMM consistently controlled type I error, while the power of LMM was lower than that of JMR-NC in most of the simulated OTUs. The ZIBR method yielded inflated type I error rate and low efficacy regardless of sample size.

**Table 1.** The type I error and power based on 10000 simulated replicates for a taxon at different levels of mean relative abundance (***y***).

| $-\log_{10}(y)$ | | (0, 1] | (1, 2] | (2, 3] | (3, 4] | (4, 5] | (5, 6] |
| --- | --- | --- | --- | --- | --- | --- | --- |
|  |  | Type I error |  |  |  |  |  |
| $N = 100$<br>($S = 50$) | JMR-NC | 0.01 | 0.003 | 0.008 | 0.0004 | 0 | 0 |
|  | JR-NC | 0.061 | 0.03 | 0.001 | 0.002 | 0.0003 | 0.0005 |
|  | LMM-N | 0.045 | 0.031 | 0.026 | 0.033 | 0.068 | 0.052 |
|  | LMM-S | 0.041 | 0.036 | 0.023 | 0.036 | 0.066 | 0.049 |
|  | ZIBR-N | 0.057 | 0.066 | 0.080 | 0.053 | 0.164 | 0.369 |
|  | ZIBR-S | 0.049 | 0.079 | 0.076 | 0.063 | 0.163 | 0.366 |
| $N = 200$<br>($S = 100$) | JMR-NC | 0.018 | 0.004 | 0.022 | 0.051 | 0.001 | 0.0003 |
|  | JR-NC | 0.162 | 0.091 | 0.049 | 0.066 | 0.035 | 0.002 |
|  | LMM-N | 0.043 | 0.039 | 0.068 | 0.057 | 0.079 | 0.058 |
|  | LMM-S | 0.046 | 0.051 | 0.066 | 0.060 | 0.079 | 0.056 |
|  | ZIBR-N | 0.056 | 0.067 | 0.099 | 0.078 | 0.170 | 0.324 |
|  | ZIBR-S | 0.059 | 0.088 | 0.099 | 0.088 | 0.174 | 0.324 |
|  |  | Power |  |  |  |  |  |
| $N = 100$<br>($S = 50$) | JMR-NC | 0.399 | 0.5 | 0.372 | 0.833 | 0.798 | 0.136 |
|  | JR-NC | 0.32 | 0.65 | 0.051 | 0.057 | 0.548 | 0.040 |
|  | LMM-N | 0.040 | 0.084 | 0.474 | 0.107 | 0.552 | 0.512 |
|  | LMM-S | 0.043 | 0.090 | 0.468 | 0.108 | 0.347 | 0.468 |
|  | ZIBR-N | 0.06 | 0.088 | 0.412 | 0.123 | 0.743 | 0.666 |
|  | ZIBR-S | 0.057 | 0.095 | 0.405 | 0.123 | 0.686 | 0.664 |
| $N = 200$<br>($S = 100$) | JMR-NC | 0.452 | 0.723 | 0.619 | 0.944 | 0.917 | 0.388 |
|  | JR-NC | 0.677 | 0.699 | 0.194 | 0.836 | 0.737 | 0.324 |
|  | LMM-N | 0.06 | 0.115 | 0.429 | 0.478 | 0.287 | 0.253 |
|  | LMM-S | 0.068 | 0.134 | 0.423 | 0.321 | 0.273 | 0.206 |
|  | ZIBR-N | 0.057 | 0.332 | 0.433 | 0.667 | 0.280 | 0.379 |
|  | ZIBR-S | 0.063 | 0.344 | 0.430 | 0.648 | 0.265 | 0.377 |

We generated 10 replicates of each OTU table to assess the performance of competing methods in scenarios B1, B2 and C. The taxa associated with disease onset in each OTU table are detected by FDR cutoff *q* < 0.15. The FPPR in scenario B1 (**Figure 3**) showed that adjusting for the top-correlated taxa in JMR successfully controlled the rate of pseudo biomarkers across distinct scenarios with power higher than LMM at larger sample size (*N* = 200), although this strategy resulted in lower detection power compared to JMR-NC. The results in **Figure 4** also demonstrated the outperformance of JMR-NC in terms of sensitivity and FDR in the taxon-specific trajectory analysis for an OTU table. The LMM methods were powerful in the test of intercept, but this model occasionally produced inflated FPPR as shown in **Figure 3** regardless of the set-level random effect size. Furthermore, the power of LMM was unstable in intercept analysis, while its power in slope analysis was nearly zero. A larger set-level random effect size yielded higher FPPR in LMM, but this impact was trivial in the other methods. JMR showed the best performance in slope analysis, with higher sensitivity (**Figure 4**) and the lowest FPPR. Results of scenario C in **Figure 5** showed that JMR and LMM effectively controlled the FPR, while LMM produced higher FPR at larger matched-set-specific random effect (*γ*). The FDR of JMR at *γ* = 0.6 was relatively higher than that of LMM due to lower sensitivity. The ZIBR methods yielded inflated FPPR in the high-dimensional scenarios, although the power of this model was higher than the competing methods regardless of sample size. The severely-inflated FPR and FDR by ZIBR in scenario C (**Figure 5**) is consistent with the results of scenario B (**Figure 3**).

**Figure 3.**
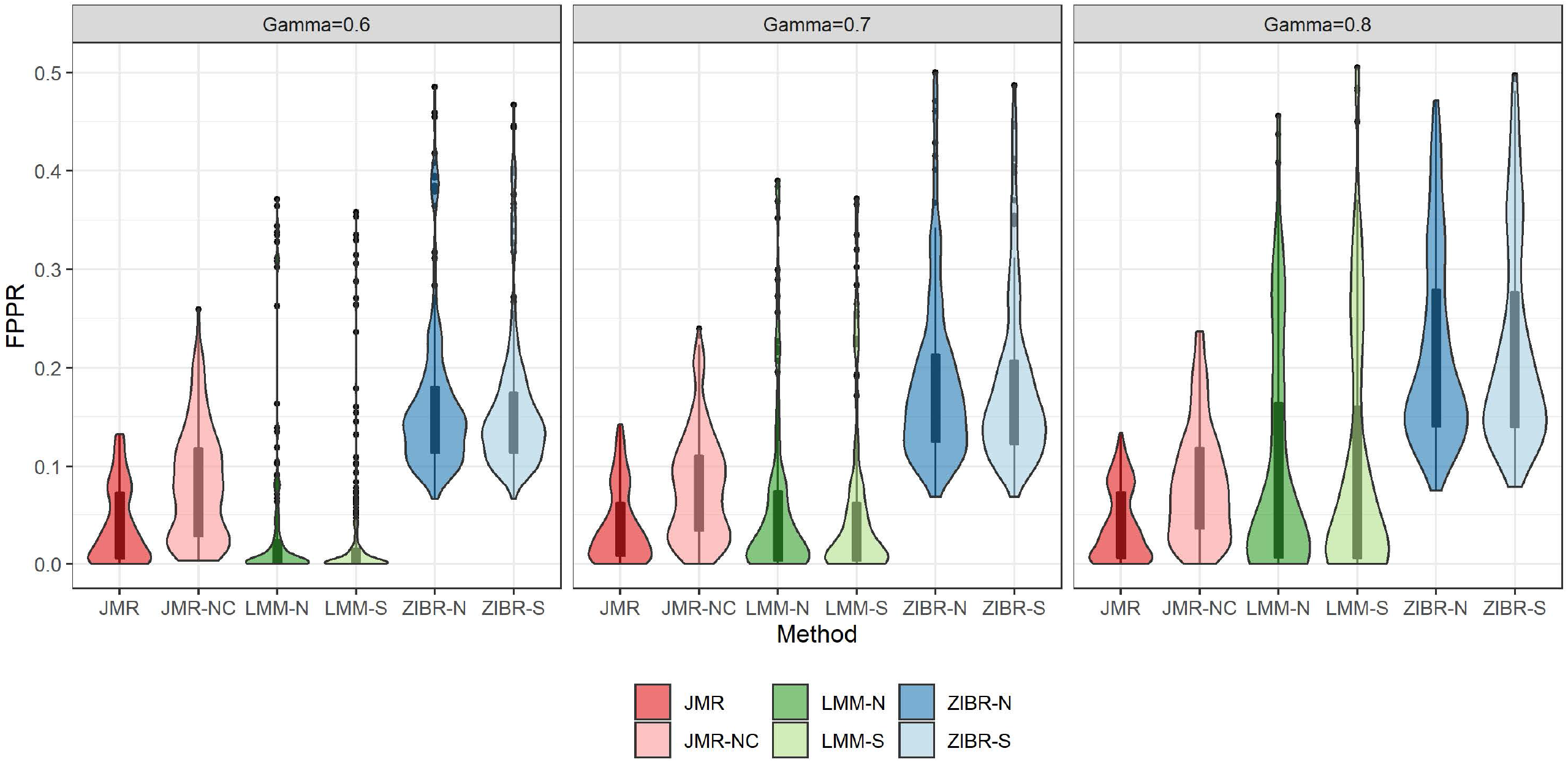
FPPR of each method for scenario B1 at different size of matched-set-specific random effect.

**Figure 4.**
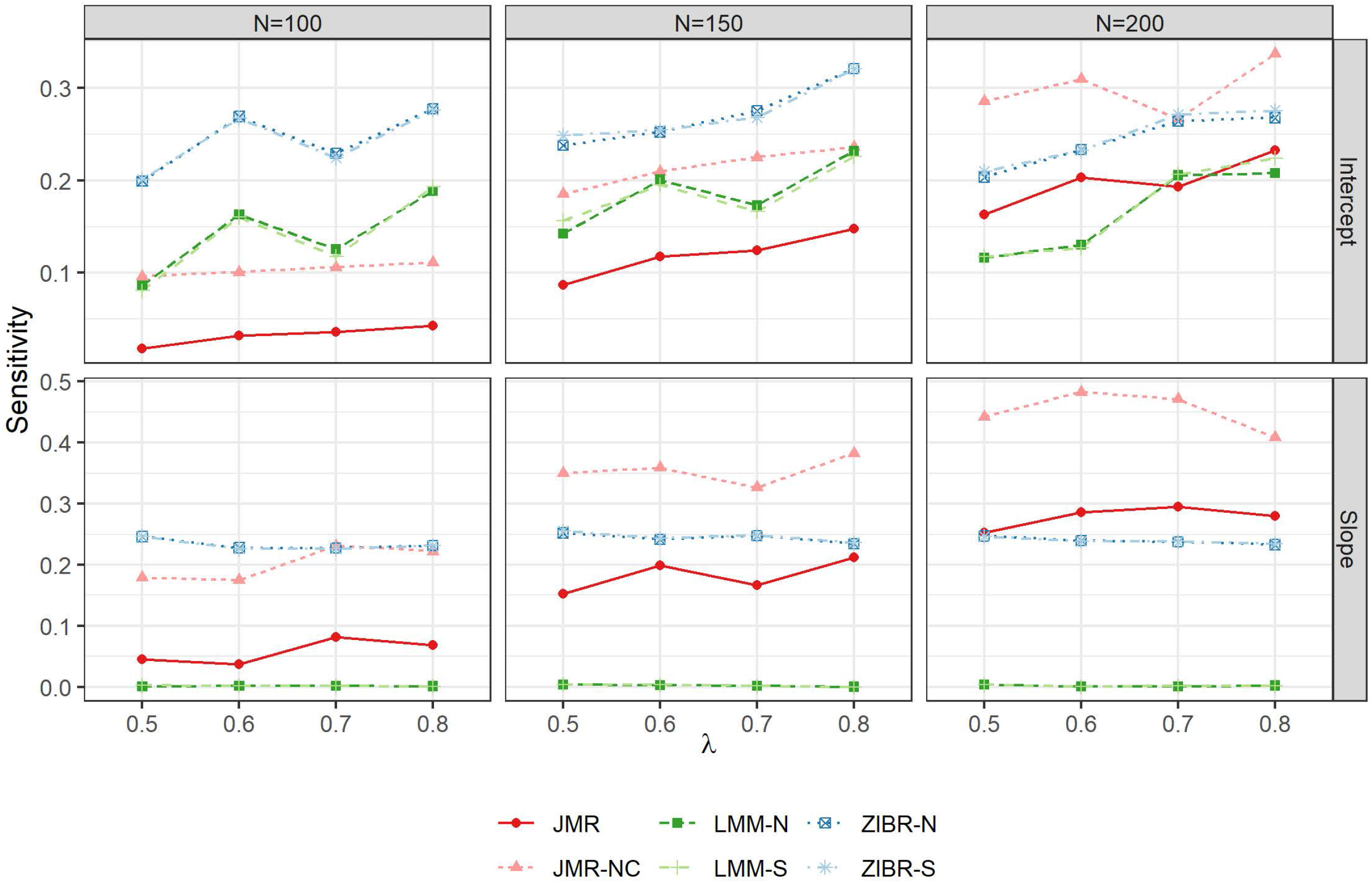
Sensitivity of each method for scenarios B1 and B2 at different sample size (*N*) and different effect size (*λ*).

**Figure 5.**
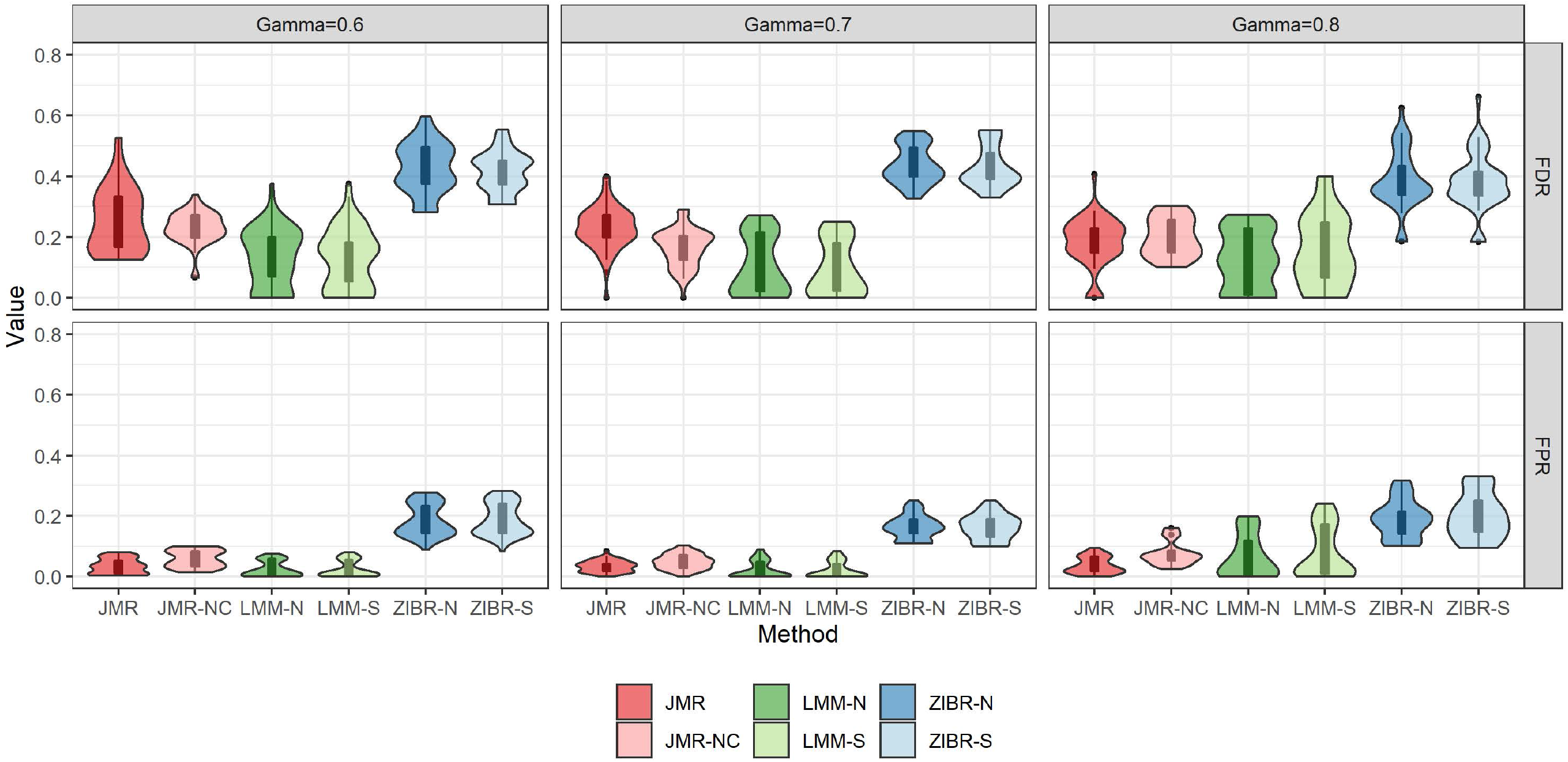
FDR and FPR for each method in scenario C at different size of matched-set-specific random effect.

### Application in TEDDY

We applied the competing methods to the longitudinal metagenomes profiled from TEDDY children’s monthly stool samples collected at the age of 4-9 months [1]. We included the cases developing IA between 9-month and 24-month age and their matched controls who remained IA-negative by the cases’ diagnosis age. For each matched pair included in the present analysis, one participant was IA positive and the other one was negative at the age of 24 months. We excluded the participant(s) matched to multiple pairs, yielding *N* = 152 subjects (*S* = 76 pairs) and *n* = 672 metagenome samples. The cases who experienced IA onset after 24 months and their matched controls were not included in this analysis.

We first filtered OTUs at genus and species level by relative abundance > 10^−6^ and prevalence > 5%, selecting 125 genera and 365 species in downstream analysis.

The sample age and the hosts’ breastfeeding status per time point were used as longitudinal covariates, while HLA DR3&4 haplotype was included as time-invariant covariates. For the LMM and ZIBR methods, we used the interaction term between IA status and binarized HLA category (DR3&4 vs. others) as a covariate to adjust for the association. We tested each OTU’s association with IA by FDR cutoff *q* <or *q* < 0.1, individually. The results in Table 2 showed that JMR identified more OTUs than LMM in both intercept and slope analysis. The LMM methods only found a small subgroup of taxa associated with IA at either genus or species level. We also visualized the overlap and difference between JMR, JMR-NC, LMM-N in **Figure 6**, and then compared Akaike Information Criterion (AIC) of JMR and JMR-NC for the 76 species detected by both methods. Adjusting for the correlated taxa in JMR did improve model fitting with lower mean AIC (−2631.847) compared to JMR-NC (−2615.571). LMM-N is not comparable to JMR or JMR-NC in terms of information criteria, since the taxon-specific relative abundance was transformed by asin-square-root.

**Table 2.** The number of genera and species associated with IA detected by each method in a subgroup of TEDDY participants.

|  |  | Intercept |  | Slope |  |
| --- | --- | --- | --- | --- | --- |
| FDR | | $q < 0.05$ | $q < 0.1$ | $q < 0.05$ | $q < 0.1$ |
| Genera | JMR | 27 | 31 | 27 | 34 |
|  | JMR-NC | 44 | 52 | 36 | 44 |
|  | LMM-N | 11 | 16 | 3 | 4 |
|  | LMM-S | 49 | 82 | 10 | 19 |
|  | ZIBR-N | 26 | 30 | 31 | 36 |
|  | ZIBR-S | 26 | 32 | 31 | 36 |
| Species | JMR | 75 | 94 | 37 | 46 |
|  | JMR-NC | 147 | 166 | 112 | 125 |
|  | LMM-N | 43 | 60 | 13 | 21 |
|  | LMM-S | 40 | 63 | 7 | 16 |
|  | ZIBR-N | 89 | 106 | 119 | 138 |
|  | ZIBR-S | 83 | 105 | 120 | 140 |

**Figure 6.**
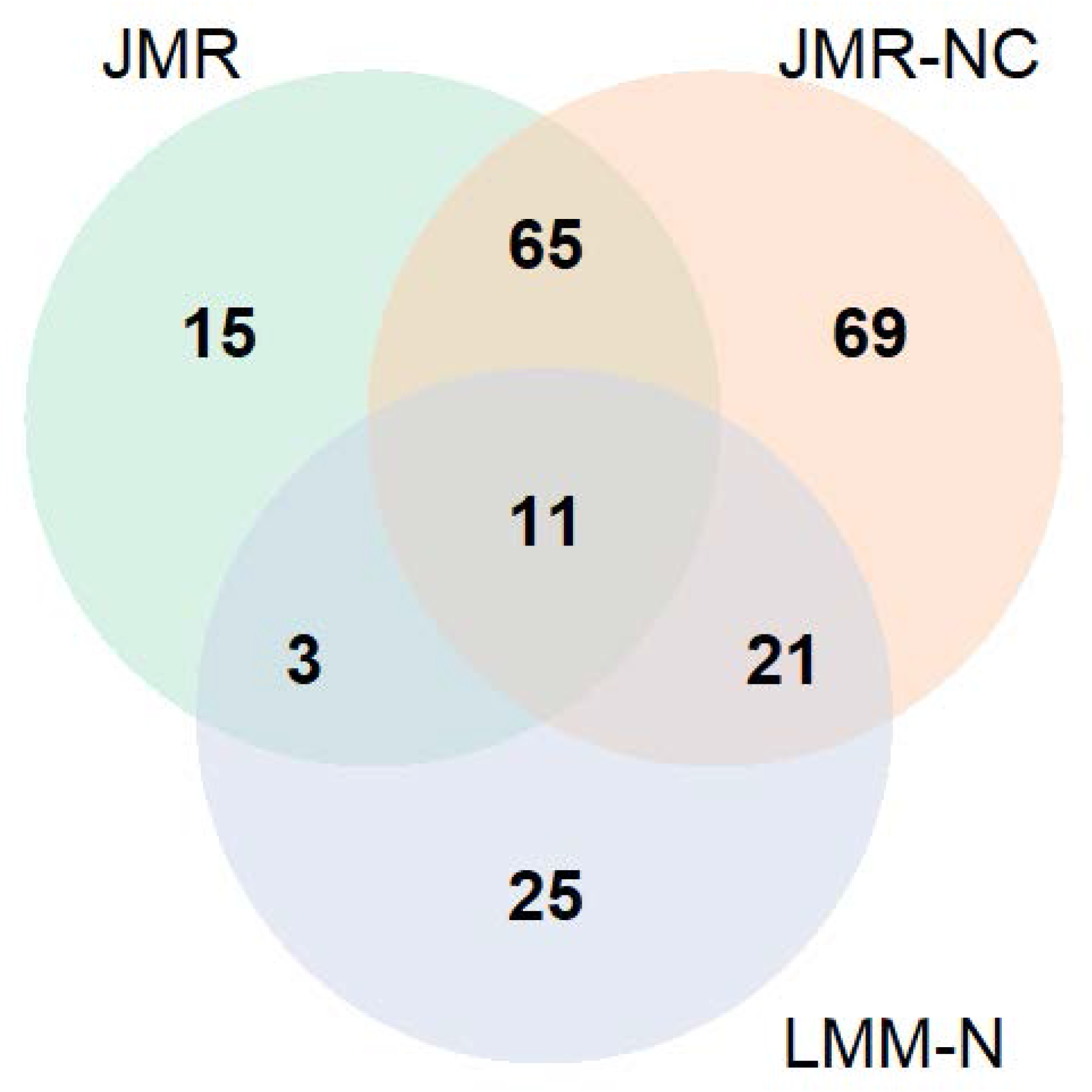
Venn diagram for the intercept analysis in TEDDY data by JMR, JMR-NC, LMM-N.

The taxa with mean abundance (intercept) associated with IA onset exclusively detected by both JMR and JMR-NC at *q* < 0.1 include *Bifidobacterium breve, Bacteroides fragilis, Lactobacillus ruminis, Veillonella ratti. B*.*breve*, as one of the three species dominating infant-age gut microbiota in TEDDY, was less abundant in intercept (i.e. at 4- and 9-month) during infancy among IA cases, with density shown in **Figure 7**. The species *B*.*fragilis* as part of the normal microbiota in human colon was found more abundant among IA cases compared to their matched controls (**Figure 1**). This *Bacteroides* species was also found differential between T1D cases and controls at only one time point in a small-size Finnish cohort [20]. There are two more abundant species *Faecalibacterium prausnitzii* and *Escherichia coli* visualized in **Figure 1** associated with IA in slope and exclusively detected by JMR. *F*.*prausnitzii*, as one of the most abundant and important commensal bacteria of human gut microbiota that produces butyrate and short-chain fatty acids from the fermentation of dietary fiber, increased faster in IA cases after 6-month of age. This rapid change and abnormally higher level of *F*.*prausnitzii* prior to IA seroconversion may be a result of the sudden change of dietary pattern during infancy.

**Figure 7.**
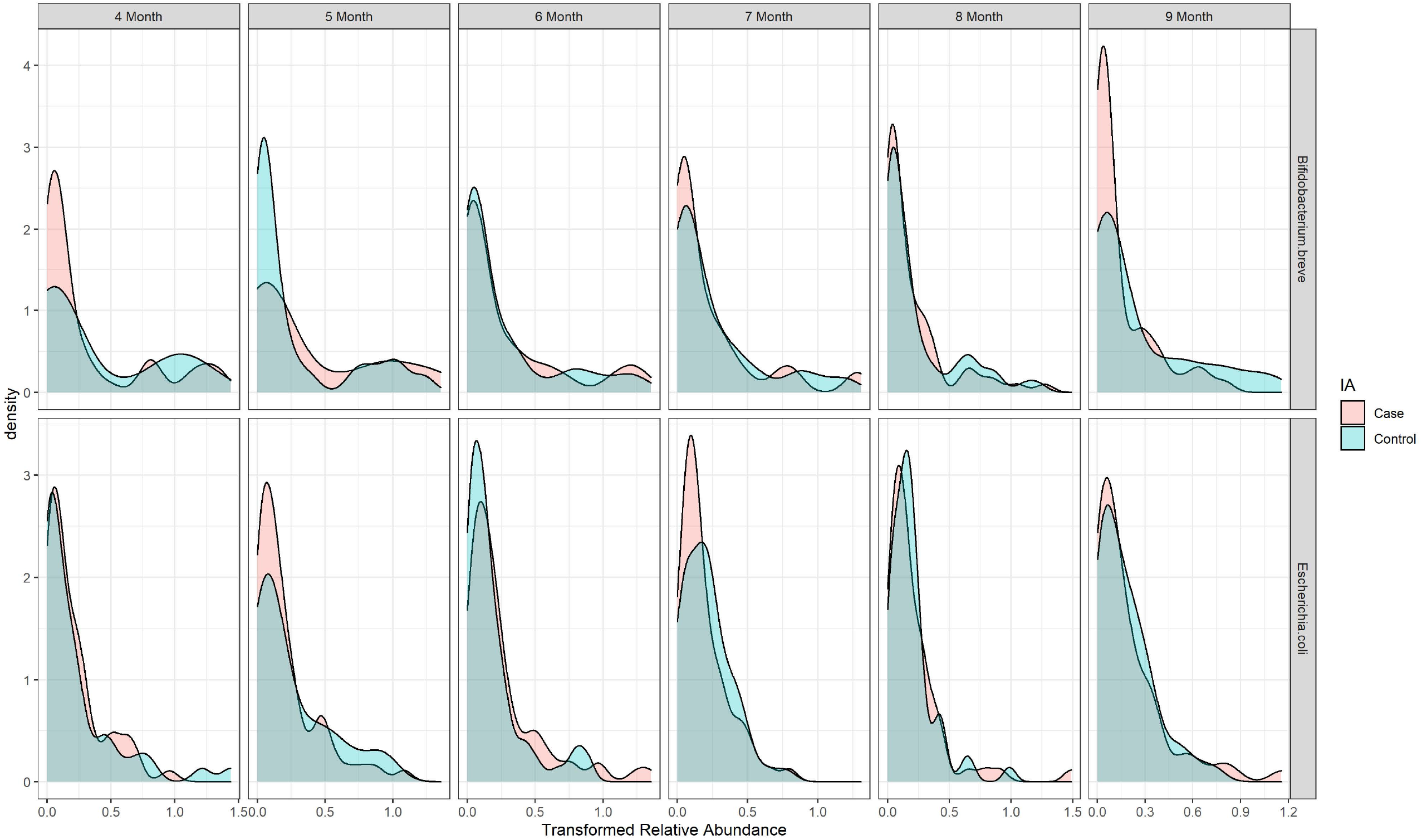
Distribution of relative abundance for *B*.*breve* and *E*.*coli* per time point grouped by 2-year IA status.

Our method successfully detected the case-control difference in the slope of *E*.*coli*, which was found as an amyloid-producing bacteria with temperal dynamics heralding IA onset in a subset analysis in DIABIMMUNE cohort [21]. The relative abundance of *E*.*coli* in TEDDY smoothly decreased from 4-month to 9-month for both cases and controls (**Figure 1**), and it was relatively more abundant in controls between 7- and 9-month with stratified densities shown in **Figure 7**. The temporal change of *E*.*coli* prior to IA seroconversion in TEDDY detected by JMR was consistent with the decrease of *E*.*coli* reported in DIABIMMUNE cohort [21], which was possibly due to prophage activation according to the *E*.*coli* phage/*E*.*coli* ratio prior to *E*.*coli* depletion in that research.

## Discussion

We developed a joint model with nested random effects to test the association between taxa and disease risk, and adjusted for the correlated taxa screened by a pre-selection procedure in abundance and prevalence, individually. We implemented our method in an R package mtradeR (metagenomic trajectory analysis with disease endpoint and risk factors), available on Github (https://github.com/qianli10000/mtradeR) and CRAN. The JMR function implemented the framework in equation (1) by parallel computing. We also provided simulation functions StatSim and TaxaSim to generate (binary) disease status and temporal high-dimensional metagenomic counts of matched sets.

The simulation of single OTU demonstrated the performance of each method at different relative abundance levels, implying that LMM with either single or nested random effect is still a robust method. The simulation of high-dimensional OTU tables also illustrated LMM’s overall performance in the test of intercept, but the unstableness of LMM is a concern in real data analysis. JMR yielded lower false or pseudo positive rate in the simulated datasets and higher detection power in slope analysis by adjusting for the top-correlated taxa in modeling. The pre-selection of top-correlated taxa in JMR was performed in relative abundance and presence, individually, being consistent with the two-part model strategy. According to the simulation study, a disadvantage of JMR is the power being relatively low at small sample size and depends on the tuning of shrinkage parameter. Adding nodes in the GH approximation may improve the power of JMR, but more nodes will also lead to additional computation costs. Hence, future work should focus on improvement of JMR in both detection power and computation efficiency.

Another limitation of our method is the occasional bias in scaling parameter (*λ*_*r*_, *λ*_*p*_) estimation, potentially caused by the *L*_2_ penalty. Our current work only focused on the unsigned association between a taxon and host disease status by using a Wald statistic. An improvement in the estimate of scaling parameter and statistical inference should be considered in future work, such as the algorithm in ZINQ [14]. We did not use quantile regression in current research, since the performance of ZINQ required tuning of grid. But ZINQ provided an alternative approach for modeling zero-inflation in microbiota composition with fewer statistical assumptions.

There are other important topics to be considered in the modeling of longitudinal microbiome data. One potential direction is high dimensional modeling framework, such as tensor singular value decomposition [22]. A promising extension of the current work in JMR is to exploit functional data analysis for multiple microbial trajectories. By employing a non-parametric joint modeling, we may be able to capture nonlinear trends and heterogeneous patterns of longitudinal biomarkers in microbiota, as well as negative correlations among taxa [23].

## Conclusions

The proposed framework JMR successfully controlled the false or pseudo biomarkers in taxon-specific trajectory analysis with improved detection power by incorporating the matching of participants and adjusting for the dependence between taxa.

## Methods

### Joint model with matching and regularization (JMR)

The probability for participant *j* (*j* = 1, …, *J*) in matched set *s* (*s* = 1, …, *S*) developing the disease of interest is 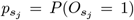, where 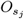 is the binary disease status. There are *J* participants in each matched set. Let 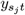 be the relative abundance of an OTU for participant *j* in matched set *s* at time point 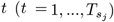. We denote the expected non-zero abundance by 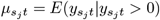, and the probability of presence (or zero-inflation) by 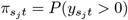, similar to [9]. For a microbiome study matching participants by the disease-associated factors and/or disease status (e.g., DIABIMMUNE, TEDDY), the matched participantsare assumed to have comparable but distinct disease risk. Hence, we model the disease status by a logistic mixed effect model with nested random effects. A joint model for the host disease status and microbial trajectory in matched set is

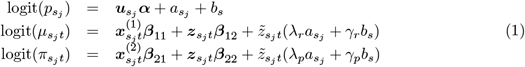

The host disease status is determined by a vector of fixed effect covariates 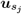 and the independent nested random effects 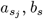. The non-zero relative abundance 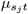 and presence 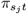 per OTU are predicted by the same random effects rescaled by parameters *λ*_*r*_, *λ*_*p*_ and a vector of clinical or bioinformatics technical covariates 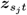. To model the unknown correlation between taxa, this OTU’s non-zero abundance and presence per time point also depend on the other taxa with relative abundance 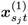 and presence-absence 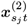 measured at the same time point, pre-selected by a procedure described below. The two-part submodel of 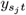 characterizes how the trajectory is affected by subject- and set-level latent factors contributing to disease risk, and how the OTU trajectory interacts with correlated taxa over time. If an OTU is a pseudo biomarker, then its relative abundance 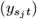 should be driven by the top-correlated taxa per time point instead of the disease-associated random effect 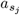. On the other hand, the abundance of a true biomarker OTU at each time point is mainly determined by the latent risk of disease onset 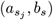 and possibly associated with the top-correlated taxa.

We set 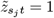 in equation (1) to test intercept, and 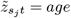 to test slope. The nested random effects and parameters *λ*_*r*_, *λ*_*p*_ provide flexibility in the modeling of between-subjects and between-sets heterogeneity, as well as model the abundance-presence correlation in each taxon by shared nested random effects instead of assuming independence between the two processes as in [9].

### Parameter estimation and hypothesis testing

To account for the sum-to-one restriction on non-zero relative abundance 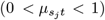 and the binarized measurement 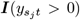 of an OTU, we intuitively employ the Zero-Inflated Beta (ZIB) density function [9] to define the match-set-specific marginal likelihood for parameter estimation. That is 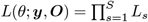, where

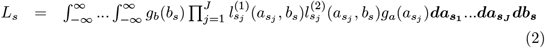

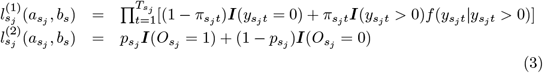

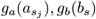 are the Gaussian density functions with mean 0 and variance 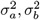, individually, and 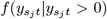 is the Beta density function with mean 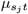 and overdispersion *ϕ*. In the simulation study, we demonstrated that the robustness and performance of this model does not require the observed relative abundance being generated from ZIB distribution.

The estimate of overdispersion 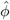 without regularization is severely inflated and also leads to bias in the estimate of other parameters. Hence, we use *L*_2_ (ridge) regularization to control the overdispersion and type I error in hypothesis testing. All the parameters *θ* are estimated by maximizing a penalized marginal likelihood function 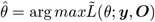, where

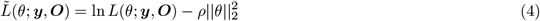

and *ρ* is selected by a cross-validation described below.

There is no closed form of the multivariate integral *L*_*s*_ in equation (2) because of the Beta density in 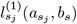. Hence, *L*_*s*_ can be approximated by Gauss-Hermite (GH) quadrature, with details explained in Appendix. We test the association between OTU trajectory and host disease status with null hypothesis:

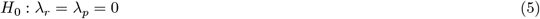

and a Wald statistic 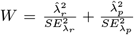, which follows a Chi-Square distribution *W* ∼ *χ*^2^(2). The false discovery rate (FDR) for multiple testing is corrected by the Benjamini-Hochberg (BH) procedure.

### Pre-selection of correlated taxa and tuning parameter selection

For each OTU 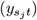 in equation (1), using all the other taxa as covariates is computationally inefficient. Hence, we use a data-driven procedure to pre-select 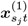 and 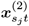, and then perform a post-selection hypothesis testing. The first step screens the taxa correlated with 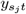 in abundance and presence, individually, using the Bray–Curtis distance [24] less than 0.1 quantile. This step may still result in many covariate taxa at species level in metageonmic data due to high dimensionality. Thus, we employ elastic net regression to further select the taxa with relative abun-dance 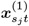 associated with 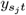 or the taxa with presence 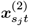associated with 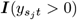, individually. In this pre-selection procedure, we model all the longitudinal metagenomes as independent samples regardless of time points (or age).

To reduce the computational burden of cross-validation for a high-dimensional OTU table, we randomly select *P*_0_ OTUs from distinct relative abundance levels to represent the complexity of microbiota composition. The matched sets are divided into 5 folds, each being a validation fold for the model built on the other four (training) folds. The penalized log likelihood in equation (4) is the negative objective function in cross-validation. For each validation fold *f* and the selected OTU *i*, the loss function is 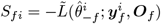, where 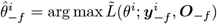. The optimal *ρ* is selected by the ‘elbow point’ minimizing 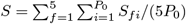.

## Availability of data and materials

The TEDDY Microbiome WGS data that supports the findings of this study have been deposited in NCBI’s database of Genotypes and Phenotypes (dbGaP) with the primary accession code phs001443.v1.p1. The R package mtradeR that implements JMR and the simulation pipeline is available at https://github.com/qianli10000/mtradeR and CRAN.

## Author’s contributions

QL proposed and implemented the model and algorithm, performed the experiments, analyzed the data, and drafted the manuscript. KV conceived the real data analysis, interpreted results, and contributed to manuscript writing. QL and YH designed the experiments. CL contributed to methodology discussion and manuscript writing. ET and LR contributed to manuscript writing. JK supervised the study design, sample collection, data generation and analysis in the TEDDY cohort.

## Acknowledgements

The work of QL was in part supported by U24DK097771 from the National Institute of Diabetes, Digestive and Kidney Diseases via the NIDDK Information Network’s (dkNET) New Investigator Pilot Program in Bioinformatics. QL and CL are in part supported by P30 Cancer Center Support Grant (CA21765) funded by National Cancer Institute; and the American Lebanese Syrian Associated Charities (ALSAC). The TEDDY study is funded by the National Institute of Diabetes and Digestive and Kidney Diseases, National Institute of Allergy and Infectious Diseases, National Institute of Child Health and Human Development, National Institute of Environmental Health Sciences, Centers for Disease Control and Prevention, and JDRF. We thank the TEDDY study data coordinating center at Health Informatics Institute, University of South Florida for data processing and sharing. We thank Suraj Sarvode Mothi for data cleaning support.

## Tables

**Additional Files**

Additional file — Appendix

