## Appendix for "A robust and transformation-free joint model with matching and regularization for metagenomic trajectory and disease onset"

For an observational study that matches participants in pairs ($J = 2$), the value of $L_s$ can be approximated by a GH quadrature

$$L_s \approx \sum_{\bar{\boldsymbol{q}}_k=(q_{1k},q_{2k},q_{3k})} \{l_{s_1}^{(1)}(q_{1k}, q_{3k}) l_{s_1}^{(2)}(q_{1k}, q_{3k})\} \times \{l_{s_2}^{(1)}(q_{2k}, q_{3k}) l_{s_2}^{(2)}(q_{2k}, q_{3k})\} \times w(\bar{\boldsymbol{q}}_k) \tag{A.1}$$

where $\bar{\boldsymbol{q}}_k$ is one of the GH nodes and $w(\bar{\boldsymbol{q}}_k)$ is the weight for each node, generated by R function mvQuad::createNIGrid. We rewrite this GH quadrature approximation in matrices below to improve computation efficiency. For each matched set, the above equation can be rewritten as

$$\begin{aligned} L_s \approx & \sum_{\bar{\boldsymbol{q}}_k=(q_{1k},q_{2k},q_{3k})} \{l_{s_1}^{(1)}(q_{1k}, q_{3k}) l_{s_2}^{(1)}(q_{2k}, q_{3k})\} \times \\ & \{l_{s_1}^{(2)}(q_{1k}, q_{3k}) l_{s_2}^{(2)}(q_{2k}, q_{3k})\} \times w(\bar{\boldsymbol{q}}_k) \end{aligned} \tag{A.2}$$

Let $l_{s_j}^{(1)}(a_{s_j}, b_s) = \prod_{t=1}^{T_{s_j}} l_{s_j t}(a_{s_j}, b_s)$, and

$$l_{s_j t}(a_{s_j}, b_s) = (1 - \pi_{s_j t}) I(y_{s_j t} = 0) + \pi_{s_j t} f(y_{s_j t} | y_{s_j t} > 0) I(y_{s_j t} > 0). \tag{A.3}$$

To calculate $L_s$ for $s = 1, \cdots, S$, we first denote

$$\bar{\boldsymbol{L}}^{(1)} = \left( \prod_{j=1}^{2} l_{1_j}^{(1)}(q_{jk}, q_{3k}), \cdots, \prod_{j=1}^{2} l_{S_j}^{(1)}(q_{jk}, q_{3k}) \right)' \tag{A.4}$$

then $\bar{\boldsymbol{L}}^{(1)}$ can be rewritten as

$$\ln \bar{\boldsymbol{L}}^{(1)} = \sum_{j=1}^{2} A_j \bar{\boldsymbol{l}}_j(\bar{\boldsymbol{q}}_k) \tag{A.5}$$

where

$$A_j = \begin{pmatrix} a_{1_j} & 0 & \cdots & 0 \\ 0 & a_{2_j} & 0 & 0 \\ \vdots & \vdots & \vdots & \vdots \\ 0 & \cdots & 0 & a_{S_j} \end{pmatrix} \tag{A.6}$$

$$a_{sj} = (1, \cdots, 1)_{1 \times T_{s_j}} \tag{A.7}$$

$$\bar{\boldsymbol{l}}_j^{(1)}(\bar{\boldsymbol{q}}_k) = \begin{pmatrix} \ln(l_{1_j 1}(q_{jk}, q_{3k})) \\ \vdots \\ \ln(l_{1_j T_{1_j}}(q_{jk}, q_{3k})) \\ \vdots \\ \ln(l_{S_j 1}(q_{jk}, q_{3k})) \\ \vdots \\ \ln(l_{S_j T_{S_j}}(q_{jk}, q_{3k})) \end{pmatrix} \tag{A.8}$$

We also denote

$$\bar{\boldsymbol{l}}^{(2)}(\bar{\boldsymbol{q}}_k) = \begin{pmatrix} \sum_{j=1}^{2} \ln(l_{1_j}^{(2)}(q_{jk}, q_{3k})) \\ \vdots \\ \sum_{j=1}^{2} \ln(l_{S_j}^{(2)}(q_{jk}, q_{3k})) \end{pmatrix} \tag{A.9}$$

Let $\bar{\boldsymbol{Q}} = (\bar{\boldsymbol{q}}_1, ..., \bar{\boldsymbol{q}}_K)$, then the matched-set-specific marginal likelihood can be approximated as

$$(L_1, \cdots, L_S)' \quad = \quad \mathbf{rowSum}\{\exp(\textstyle\sum_{j=1}^{2} A_j \bar{\boldsymbol{l}}_j^{(1)}(\bar{\boldsymbol{Q}}) + \bar{\boldsymbol{l}}^{(2)}(\bar{\boldsymbol{Q}})) \odot w(\bar{\boldsymbol{Q}})\} \tag{A.10}$$

where $\odot$ represents element-wise multiplication of matrices and **rowSum** is the row-wise sum of elements in a matrix.
